# Sustained exposure to CAR-T cell secretome impairs human Hematopoietic Stem Cell function and is reversible by dual TNFα-IFNγ blockade

**DOI:** 10.64898/2026.03.17.712280

**Authors:** Siva Sai Naga Anurag Muddineni, Diana Rasoulouniriana, Amilia Meir, Danielle Geller, Debanjan Singha Roy, Einat Tako, Neta Solomon, Tal Avraham, Yael Raz, Rony Chen, Eric Shifrut, Elad Jacoby, Michael Milyavsky

## Abstract

Prolonged cytopenias are a frequent complication of chimeric antigen receptor (CAR) T-cell therapies and are associated with increased infection risk and non-relapse mortality. Although inflammatory cytokines released during CAR-T cell activation have been implicated in immune effector cell-associated hematotoxicity (ICAHT), their direct effects on human hematopoietic stem and progenitor cells’ (HSPCs) function remains incompletely understood. Here, we established a reductionist model of CAR-T-associated hematotoxicity using conditioned media (CM) derived from activated CD19 CAR-T cells. Sustained exposure of human HSPCs to CAR-T-derived inflammatory secretome impaired HSPC expansion and reduced long-term repopulating capacity in xenotransplantation assays. In contrast, short-term exposure did not abrogate HSPC function, indicating that brief inflammatory signals can initiate durable reprogramming events, with functional consequences emerging during subsequent proliferative expansion. Mechanistically, CAR-T CM induced IFNγ- and TNFα-responsive transcriptional programs in HSPCs and promoted inflammatory myeloid skewing without evidence of apoptosis-dependent stem cell loss. Combined inhibition of IFNγ and TNFα restored HSPC expansion, normalized lineage output, reversed inflammatory transcriptional signatures, and rescued *in vivo* repopulating capacity without impairing CAR-T cytotoxic activity. These findings demonstrate that CAR-T-derived inflammatory signaling can directly impair human HSC function and identify dual TNFα/ IFNγ blockade as a potential strategy to mitigate CAR-T-associated hematotoxicity while preserving antitumor efficacy.

---

Chimeric antigen receptor (CAR) T-cell therapies have transformed treatment outcomes in a range of B cell malignancies and are increasingly explored for additional cancers and non-malignant diseases^1^. Hematotoxicity, namely post-CAR neutropenia and thrombocytopenia, has emerged as a significant complication following CD19 and BCMA directed CAR-T therapies and may persist well beyond day 30^2^. Prolonged cytopenias are associated with transfusion dependence, extended hospitalization, increased risk of severe infections, which is the leading cause of non-relapse mortality following CAR-T therapy^3^.

Accumulating evidence implicates CAR-T-derived inflammatory damage to Hematopoietic Stem and Progenitor Cells (HSPCs) in driving prolonged cytopenias, now collectively termed as immune effector cell associated hematotoxicity (ICAHT)^3–6^. Although inflammatory cytokines that are abnormally upregulated in CAR-T–treated patients’ bone marrow (BM) (e.g., IFNγ, TNFα, IL-6 and IL-1β) have been individually implicated in HSPC dysfunction^7–9^, their collective impact on human HSPC functionality remains unclear. Modeling of ICAHT has been mainly limited to transcriptomic analysis of samples from patients, reporting inflammation-related transcriptional signatures in HSPCs^10,11^. The direct effectors and functional consequences of CAR-mediated inflammation on human HSPCs remain poorly characterized, limiting the development of effective mechanism-based strategies to mitigate hematotoxicity.

To address these gaps, we established a reductionist model of CAR-T–associated hematotoxicity (Fig. 1A). Conditioned medium (CM) was collected after co-culturing human CD19 CAR-T cells with B-ALL (CAR-T CM) or untransduced T-cells with B-ALL (UT CM). Human cord-blood derived CD34+ HSPCs were exposed to UT CM or CAR-T CM for 1, 3, or 7 days, followed by functional repopulating assays and complementary immunophenotypic analysis (Fig. 1A).

**Figure 1:**
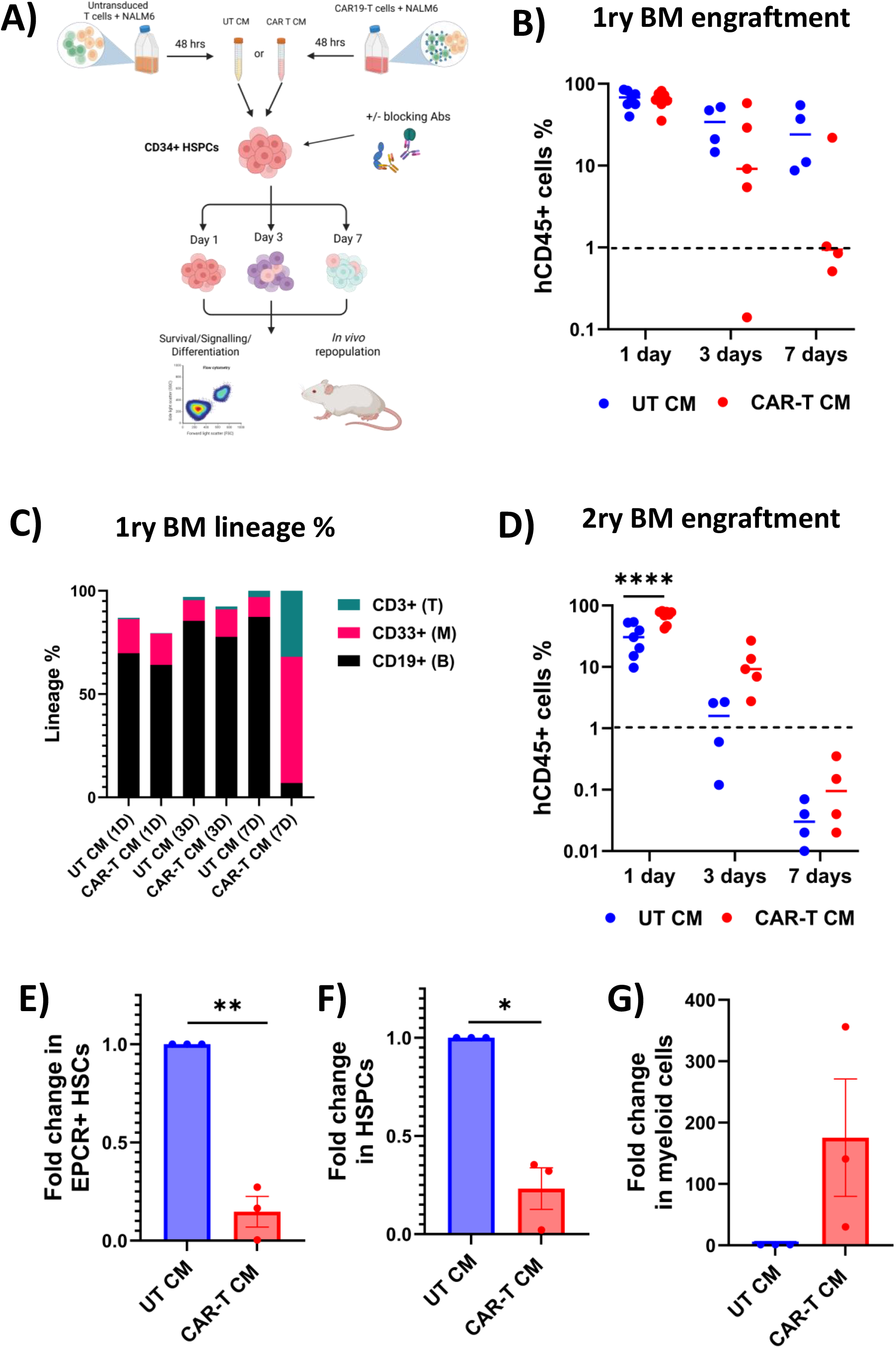
Chronic CAR-T secretome depletes HSC function by inducing myeloid differentiation. **(A)** Experimental schematic. Conditioned medium (CM) was generated from 48-hour co-cultures of untransduced (UT) T cells or CD19 CAR-T cells with NALM6 target cells. Human CD34+ HSPCs were exposed to UT CM or CAR-T CM for 1, 3, or 7 days, followed by phenotypic analysis by flow cytometry and functional assessment by xenotransplantation into NSG mice. **(B)** Primary bone marrow engraftment at 12 weeks following transplantation of CD34+ cells exposed to UT CM or CAR-T CM for 1, 3, or 7 days. Human CD45+ (hCD45+) cells are shown as percentage of total bone marrow cells. For the 1-day time point, n = 8 mice per group (two independent experiments). For the 3-day and 7-day time points, n = 4 mice per group (single experiment). **(C)** Lineage distribution in primary bone marrow at 12 weeks (CD19+ B cells, CD33+ myeloid cells, CD3+ T cells). **(D)** Secondary bone marrow engraftment at 12 weeks following transplantation of bone marrow mononuclear cells harvested from primary recipients. hCD45+ engraftment is shown for 1-day, 3-day, and 7-day exposure groups. **(E–G)** Fold change in absolute cell numbers relative to UT CM controls from sorted EPCR+CD34+CD33+CD45RA− cells (7 days exposure): (C)EPCR+ HSC fraction (EPCR+CD34+CD33+CD45RA− gate), (D) HSPC fraction (CD34+CD33+CD45RA− gate), and (E) CD11b+ myeloid cells (CD34−CD33+CD11b+ gate). Data are presented as mean ± SEM unless otherwise indicated. For engraftment studies, individual mice and median values are shown. Statistical analysis was performed using unpaired Welch’s t test (two-group comparisons) or one-way ANOVA (multiple comparisons).

We first assessed long-term multilineage engraftment potential of human HSPCs following transplantation of CD34+ cells exposed to UT CM or CAR-T CM. While short-term (1-3 day) exposure of HSPCs to either CM revealed comparable BM, spleen and peripheral blood (PB) chimerism 12-weeks post transplantation, prolonged exposure to CAR-T CM resulted in severely impaired human engraftment (Fig. 1B; Supplemental Fig. 1A–B). Here, engrafted cells displayed reduced B-cell output with relative predominance of myeloid and T-cell lineages (Fig. 1C). In serial transplantation assays –short-term exposure of CD34+ cells to CAR-T CM produced a two-fold higher human chimerism compared with UT CM controls, indicating maintenance of long-term repopulating HSPCs (Fig. 1D; Supplemental Fig.1C-D). On the other hand, HSPCs exposed to CAR-T CM for seven days failed to generate secondary grafts.

To explore potential cellular mechanisms underlying these functional alterations, we performed immunophenotypic analysis of HSPCs following CAR-T CM exposure. Within 24 hours of exposure, we observed a marked reduction in the CD34+CD38− fraction (Supplementary Fig. 2A). This change seems to reflect rapid induction of CD38 on the cell surface of CD34+CD45RA− cells, and was primarily mediated by IFNγ (Supplementary Fig. 2B–E), similar to previous reports in AML blasts^12^. Accordingly, when we measured frequency of EPCR+CD34+CD45RA-HSCs, it remained stable at 24 hours (Supplementary Fig. 3A-B). These findings indicate that inflammatory stimulation induces CD38 expression on primitive HSPCs, rendering CD38 an unreliable marker for identifying human HSCs under inflammatory conditions.

To determine the impact of sustained inflammatory exposure on primitive HSCs, we cultured sorted CD34+CD38−CD45RA−EPCR+ HSCs in UT CM or CAR-T CM for 7 days. CAR-T CM reduced HSC expansion approximately five-fold and HSPC expansion four-fold while markedly increasing CD11b+ myeloid output and inflammatory CD38 induction (Fig. 1D– F; Supplementary Fig. 3C–H). Similar trends were observed using bulk CD34+ HSPCs (Supplementary Fig. 4A–H). Notably, transient exposure to CAR-T CM for only 3 hours was sufficient to induce expansion defects and myeloid skewing despite normalization of acute inflammatory markers. Pharmacologic inhibition of apoptosis or necroptosis failed to rescue expansion, indicating that CAR-T CM primarily drives HSPC differentiation and impaired self-renewal rather than cell death.

Several inflammatory cytokines produced by activated CAR-T cells have been implicated for mediating CAR-T–associated hematotoxicity^13^. In murine models, IFNγ and IL6 have been linked to HSC dysfunction during CAR-T–induced systemic inflammation^14^, whereas single-cell analyses of patient CD34+ cells following CAR-T therapy suggest TNFα as a dominant cytotoxic mediator^11^. To dissect the direct contribution of these cytokines in our human HSPC system, we inhibited IFNγ using a neutralizing antibody, TNFα using the TNF receptor–Fc fusion protein (etanercept), and IL-6 using the IL-6 receptor antagonist (LMT-28). Neither IFNγ nor TNFα blockade alone restored overall HSPC expansion. However, combined IFNγ and TNFα inhibition significantly rescued HSPC numbers after 7 days (Fig. 2A, Supplemental Fig. 5C). In addition, TNFα blockade alone was sufficient to reverse CAR-T CM–induced CD11b+ myeloid expansion (Fig. 2B), indicating a dominant role for TNFα in promoting inflammatory myeloid differentiation. Combined blockade also led to the rescue of primitive EPCR+ HSC expansion in the presence of CAR-T CM (Supplementary Fig. 5A–B). IL6-blockade had no impact on either compartment. Accordingly, we detected significantly higher levels of IFNγ and TNFα in CAR-T CM compared to UT CM (Fig. 2C–D).

**Figure 2:**
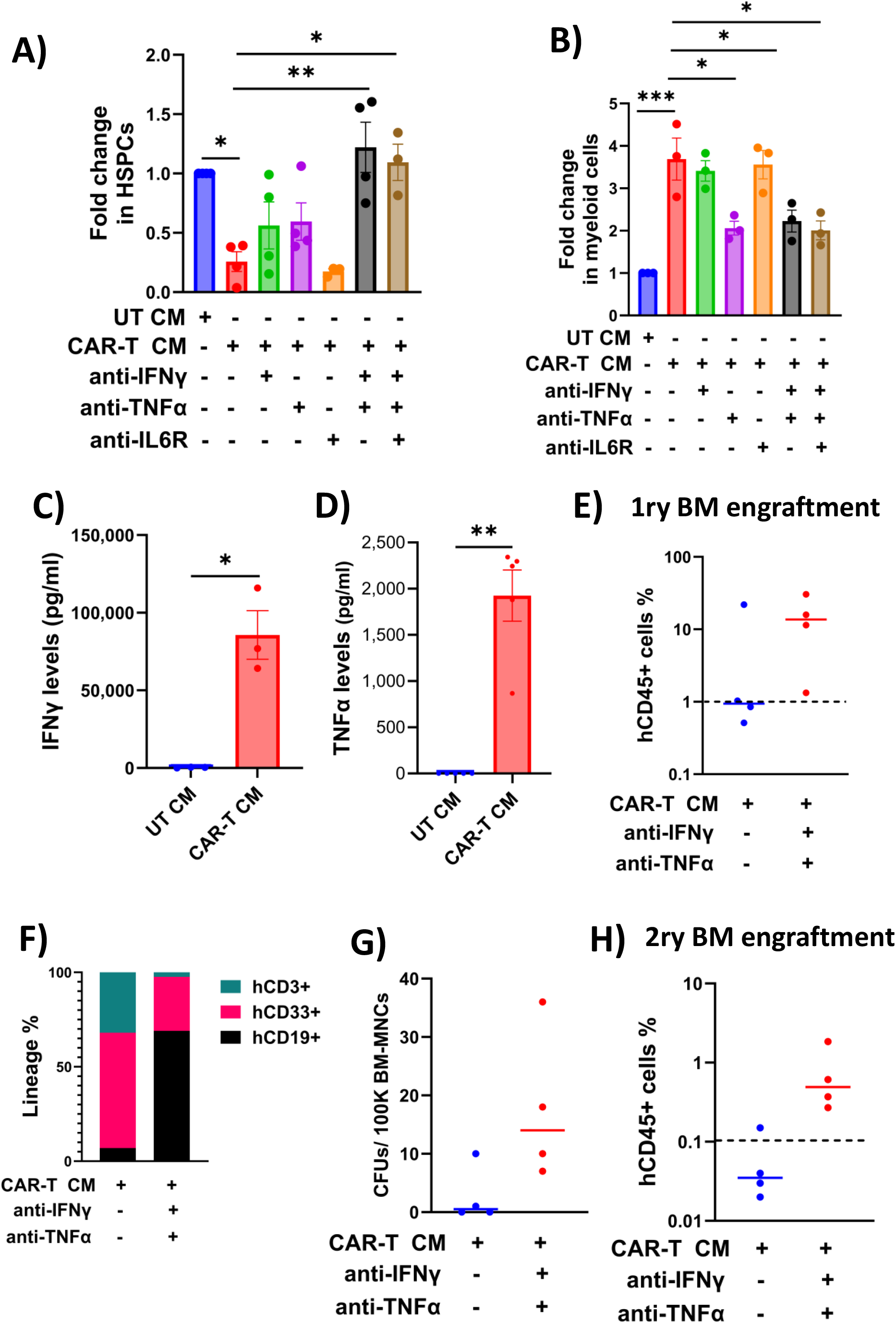
TNFa and IFNg mediate CAR-T CM induced myeloid skewing. **(A–B)** CD34+ HSPCs were cultured for 7 days in UT CM or CAR-T CM in the presence of neutralizing antibodies against IFNγ, TNFα and IL6 receptor antagonist (LMT-28), or their combinations. (A) Fold change in expansion of CD34+CD33+CD45RA− HSPCs relative to UT CM controls. (B) Fold change in CD34−CD33+CD11b+ myeloid cell expansion relative to UT CM controls. **(C–D)** Quantification of IFNγ (C) and TNFα (D) concentrations in UT CM and CAR-T CM by ELISA. Cytokine levels are shown prior to dilution for HSPC culture. **(E)** Primary bone marrow engraftment at 12 weeks following transplantation of CD34+ cells exposed for 7 days to UT CM, CAR-T CM, or CAR-T CM with combined IFNγ and TNFα blockade. Human CD45+ (hCD45+) engraftment is shown as percentage of total bone marrow cells (n = 4 mice per group; single experiment; NSGW41 recipients). **(F)** Lineage distribution of human grafts in primary bone marrow at 12 weeks, showing proportions of CD19+ B cells, CD33+ myeloid cells, and CD3+ T cells. **(G)** Colony-forming unit (CFU) assay from primary bone marrow mononuclear cells, expressed as colonies per 100,000 BM-MNCs. **(H)** Secondary bone marrow engraftment at 12 weeks following transplantation of bone marrow mononuclear cells harvested from primary recipients (n = 4 mice per group; NSGW41 recipients). Data are presented as mean ± SEM unless otherwise indicated. For engraftment studies, individual mice and median values are shown. Statistical analysis was performed using unpaired Welch’s t test (two-group comparisons) or one-way ANOVA (multiple comparisons).

To functionally asses the effects of cytokine inhibition on HSPC function, CD34+ cells were cultured with CAR-T CM for 7 days with or without cytokine blockade prior to transplantation. While only 1/4 mice engrafted following CAR-T CM exposure, combined IFNγ/TNFα blockade restored durable engraftment in 3/4 mice (Fig. 2E). Bone marrow lineage distribution normalized from myeloid-biased profile toward a more balanced lymphoid–myeloid output in the presence of IFNγ/TNFα blockade (Fig. 2F). Cytokine blockade also restored immature myeloid progenitors in bone marrow, consistent with preservation of functional HSPCs. (Fig. 2G). Secondary transplantation further confirmed functional rescue of HSPCs: whereas CAR-T CM–exposed cells failed to generate secondary grafts (0/4), combined cytokine blockade sustained engraftment in 4/4 recipients (Fig. 2H). Peripheral blood and spleen engraftment showed similar trends (Supplementary Fig. 6A–D). Importantly, IFNγ/TNFα blockade did not compromise CAR-T cytotoxic function in a 24-hour killing assay against Nalm6 (Supplementary Fig. 6E).

To further substantiate the role of IFNγ and TNFα in CAR-T–mediated hematotoxicity, we characterized transcriptional responses in CD34+ HSPCs after 3 and 24 hours of exposure. CAR-T CM induced canonical TNFα- and IFNγ-responsive genes, including IFIT1–3, BATF family members, IRF1, STAT1, NFKBIA, TRAIL, CCL2, and IL6 (Supplementary Fig. 7A–B). Although early (3-hour) induction was only partially attenuated by cytokine blockade, combined IFNγ/TNFα inhibition normalized inflammatory gene expression by 24 hours to levels comparable to UT CM controls.

Collectively, our findings demonstrate that CAR-T–derived inflammatory secretome directly impairs human HSC function. Our data support a model in which CAR-T– associated inflammation rapidly reprograms HSPCs toward precocious differentiation, with defects becoming apparent during subsequent proliferative expansion. Although overt myeloid skewing is not observed clinically in CAR-T–associated cytopenias, inflammatory activation of differentiation programs may represent an early event preceding marrow suppression^15^. Clinical manifestations may be further shaped by lymphodepleting chemotherapy, infections, antibiotic exposure, and diseased marrow microenvironments.

Our findings identify inflammatory cytokine signaling as a tractable therapeutic axis in ICAHT. Dual TNFα and IFNγ blockade restored HSPC expansion, normalized lineage output, reversed inflammatory transcriptional programs, and rescued *in vivo* repopulating capacity without compromising CAR-T cytotoxicity. Consistent with prior studies showing that IFNγ inhibition can mitigate CAR-T toxicity while preserving anti-tumor efficacy^16,17^, our data suggest that combined targeting of IFNγ and TNFα may represent a clinically actionable strategy to preserve HSC function and mitigate prolonged cytopenias following CAR-T therapy.

## Acknowledgments

This work was partially supported by Israel Science Foundation (ISF 3480/19, ISF 1362/20), Sagol Center for Regenerative Medicine and Varda and Boaz Dotan Research Center in Hemato-Oncology grants (to MM). This work was performed in partial fulfillment of the requirements for PhD degree of SSNAM, the Dr. Miriam and Sheldon G. Adelson Graduate School of Medicine, Gray Faculty of Medical and Health Sciences, Tel Aviv University, Israel.

S.S.N.A.M. received travel grants from the European Hematology Association (EHA), European Society for Blood and Marrow Transplantation (EBMT) and TAU Constantiner Institute of Molecular Genetics related to this work. The authors would like to thank Prof. Motti Gerlic (TAU) for providing Necroptosis and NLRP3 inhibitors. The authors would like to thank Dr. Irena Shur, Dr. Daria Makarovsky and Ms. Dana Bistriz of the Research Infrastructure Core Facilities at the Gray Faculty of Medical & Health Sciences of Tel-Aviv University for their technical assistance.

## Author Contributions

SSNAM, EJ and MM conceptualized the study, designed and analyzed experiments. SSNAM, DR, AM, DG and DSR performed experiments; ET, YR, TA, RC, ES, NS prepared and provided study material for experiments; SSNAM, EJ and MM wrote the original manuscript; MM provided financial support.

## Ethics statement

The study design was approved by the Institutional Review Boards of Tel Aviv University (Approval no. 0006957-5) in accordance with the relevant guidelines and regulations. Informed consent was obtained from all participants for the use of their cord blood for the isolation of CD34+ cells used in this study. All animal experimental protocols were approved by the Institutional Animal Care and Use Committee of Tel-Aviv University, Israel (TAU-MD-IL-2307-152-5).

## Data availability

Not applicable

---

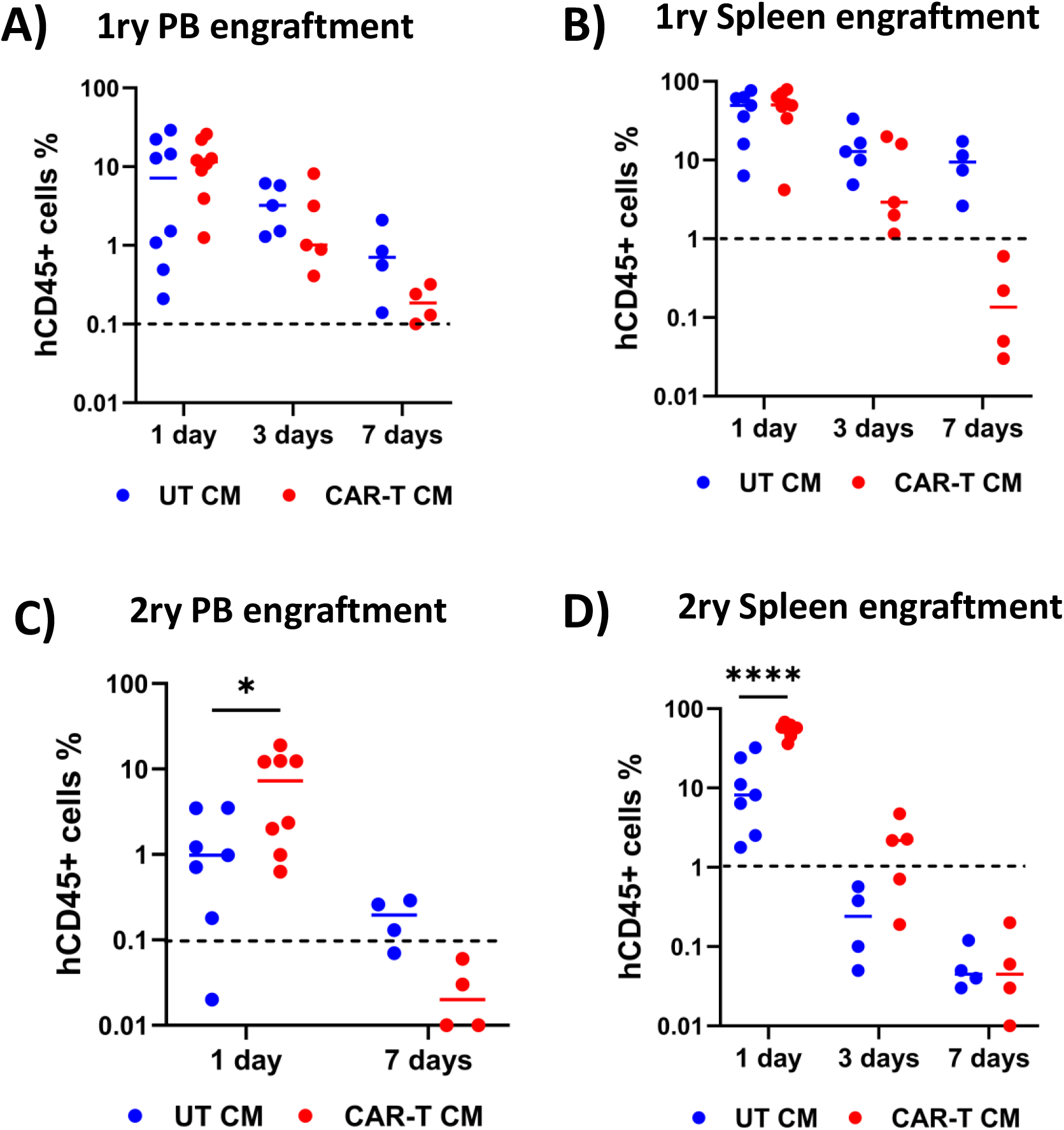

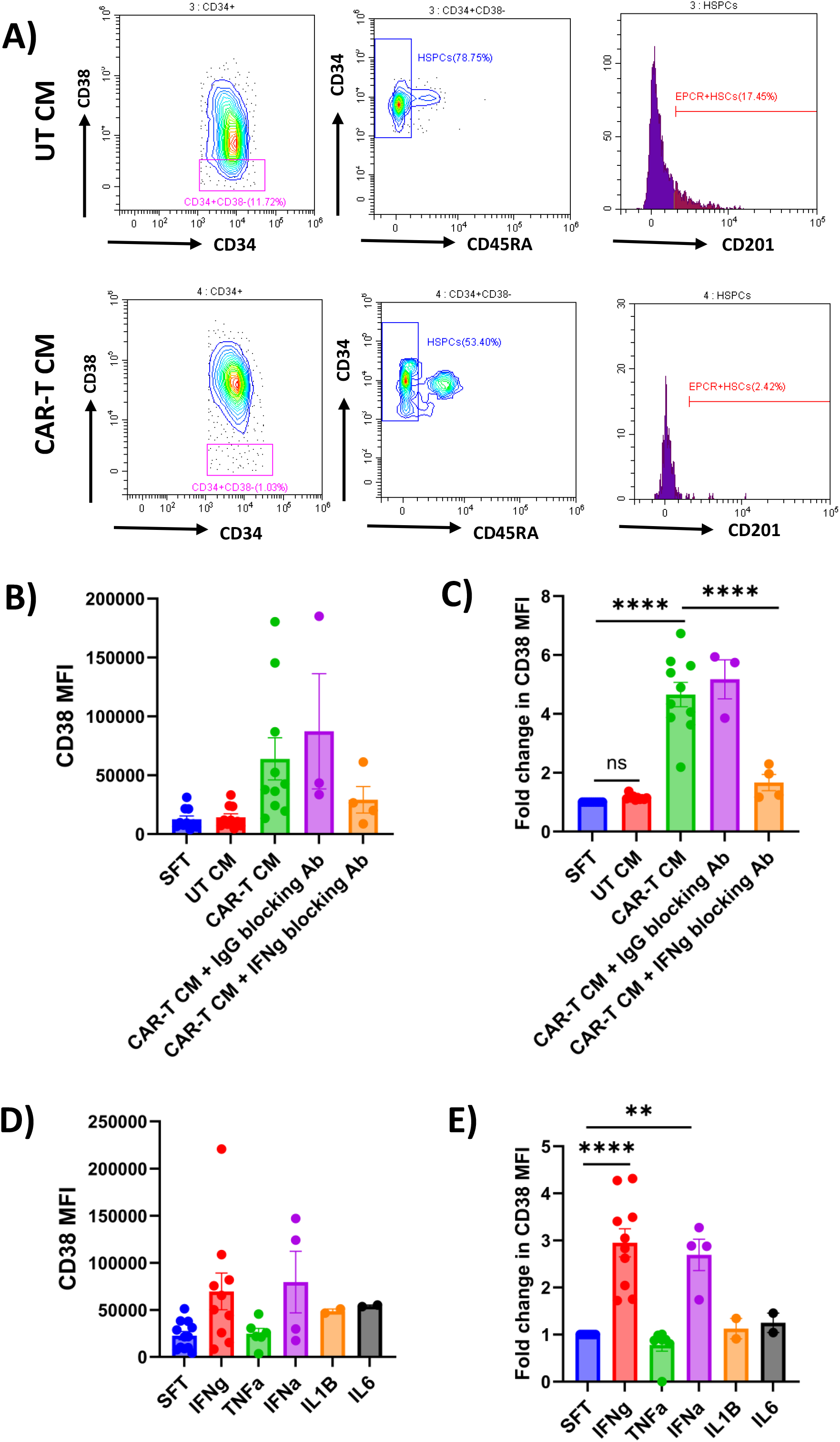

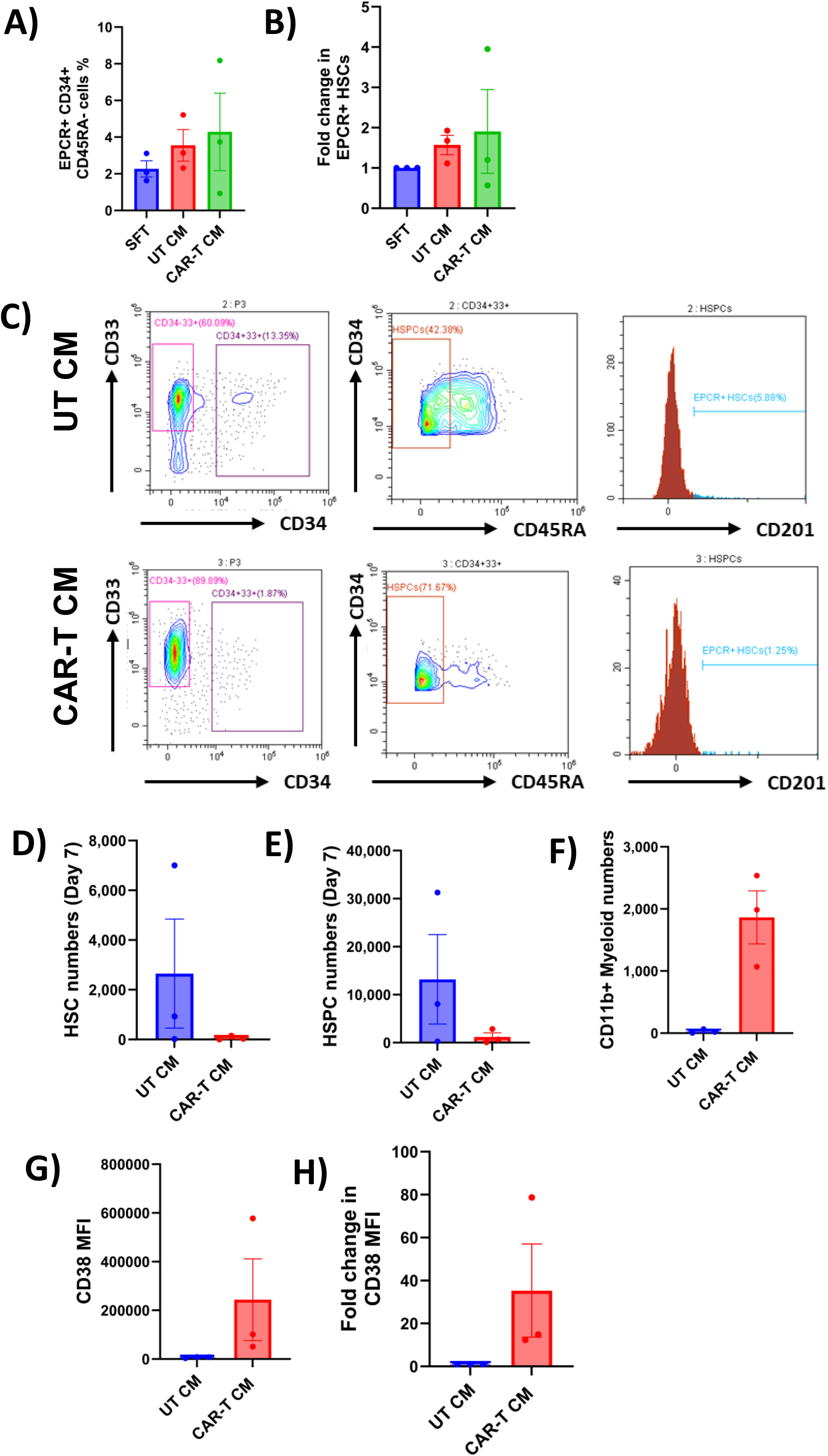

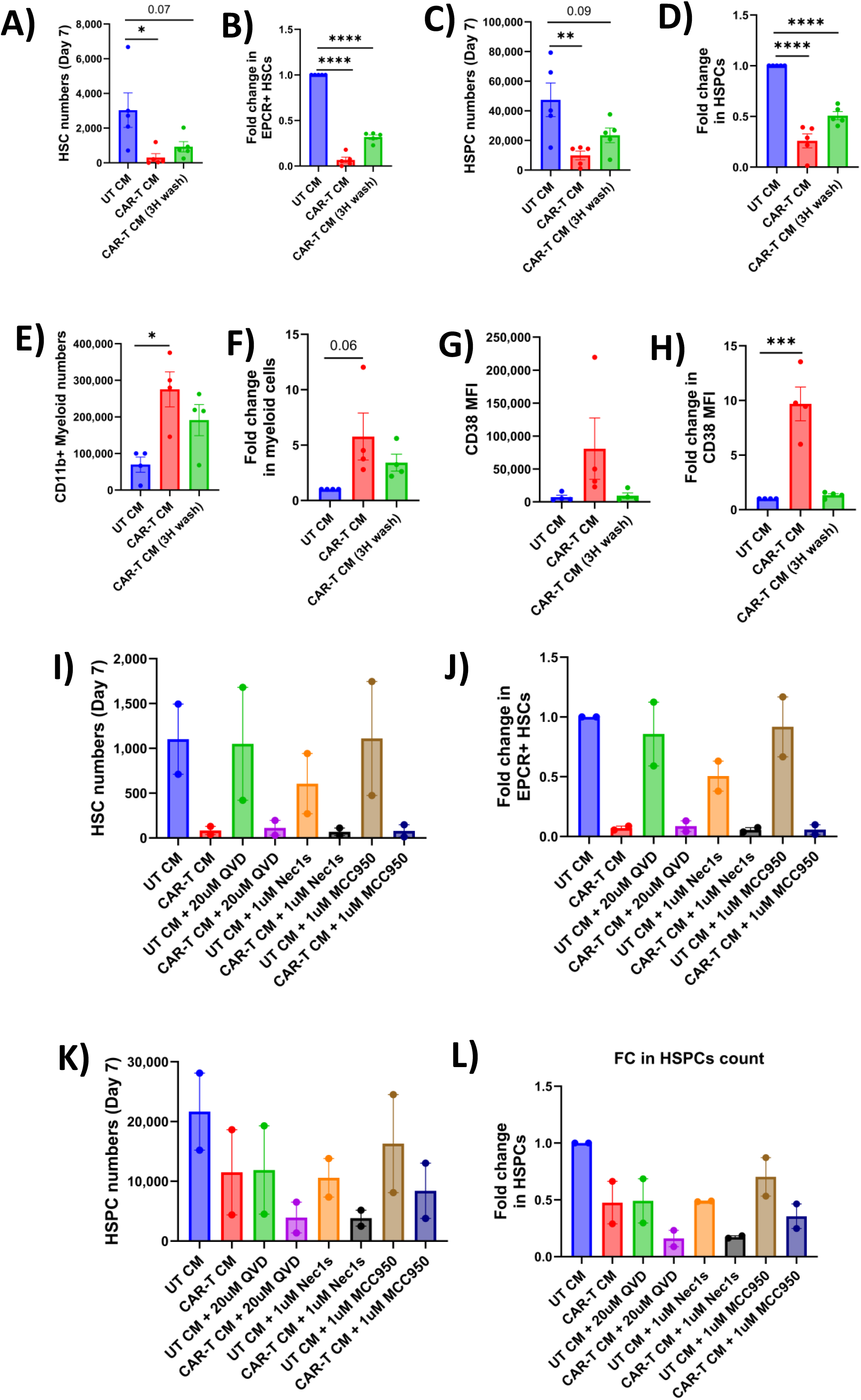

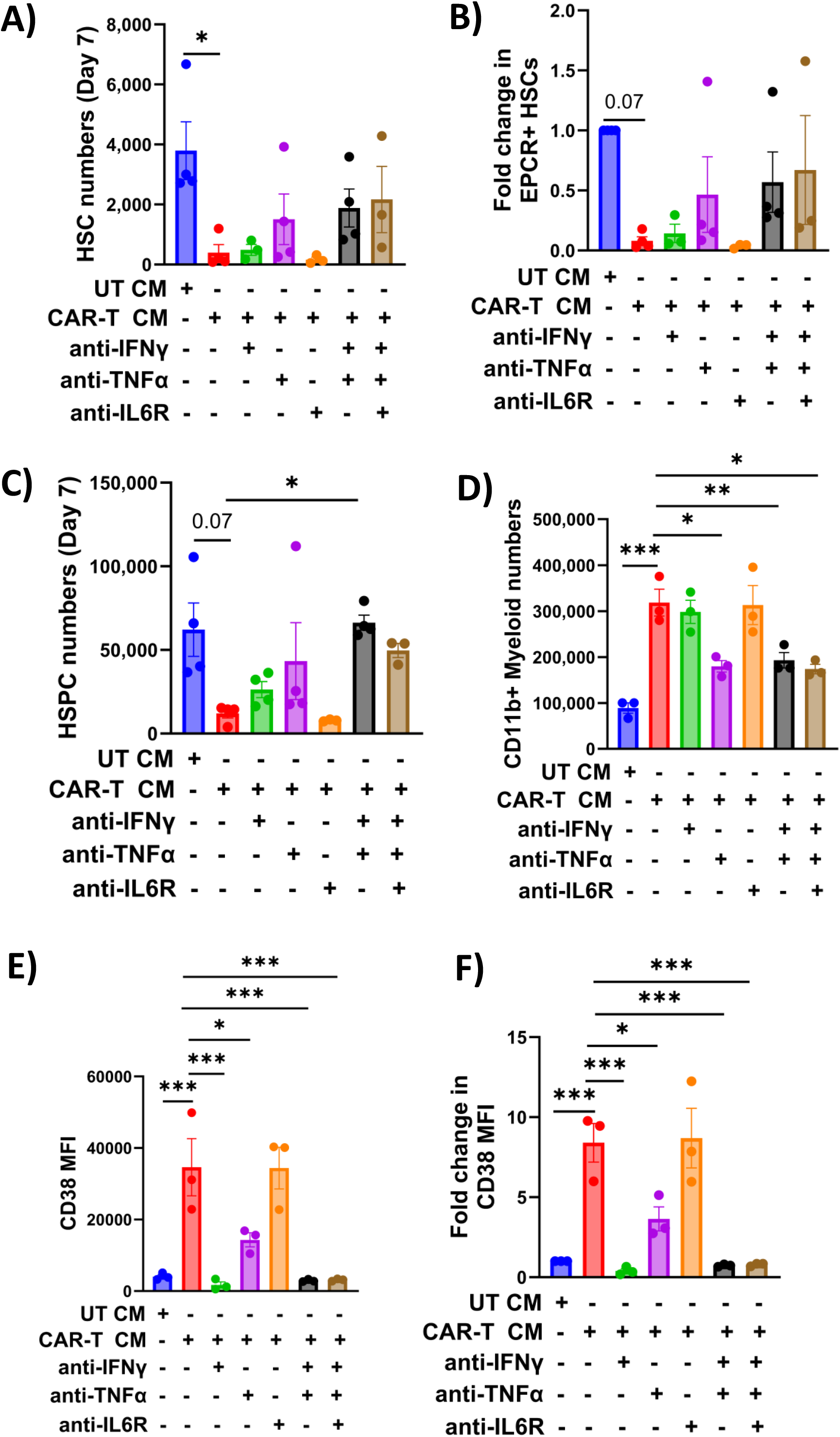

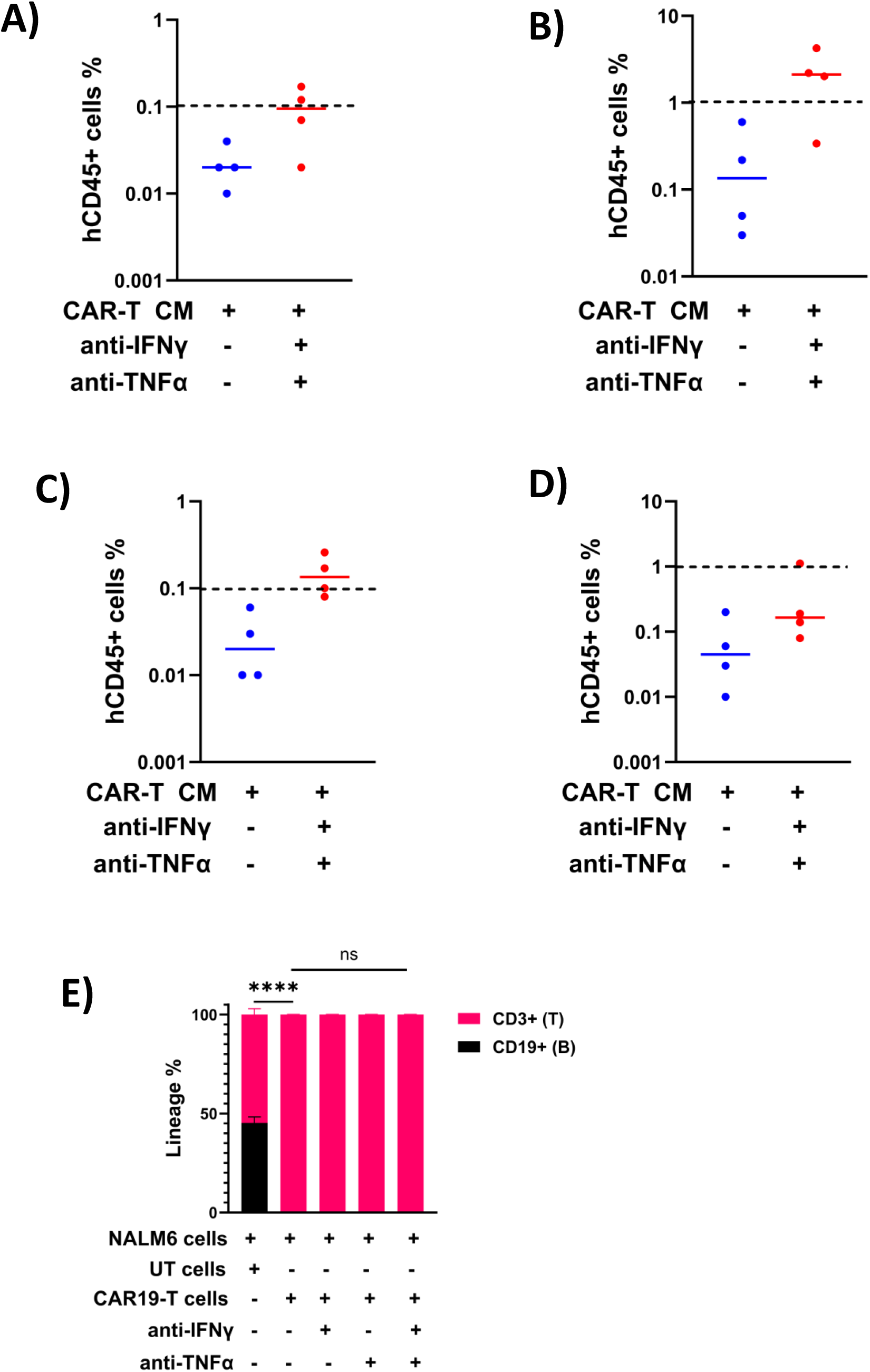

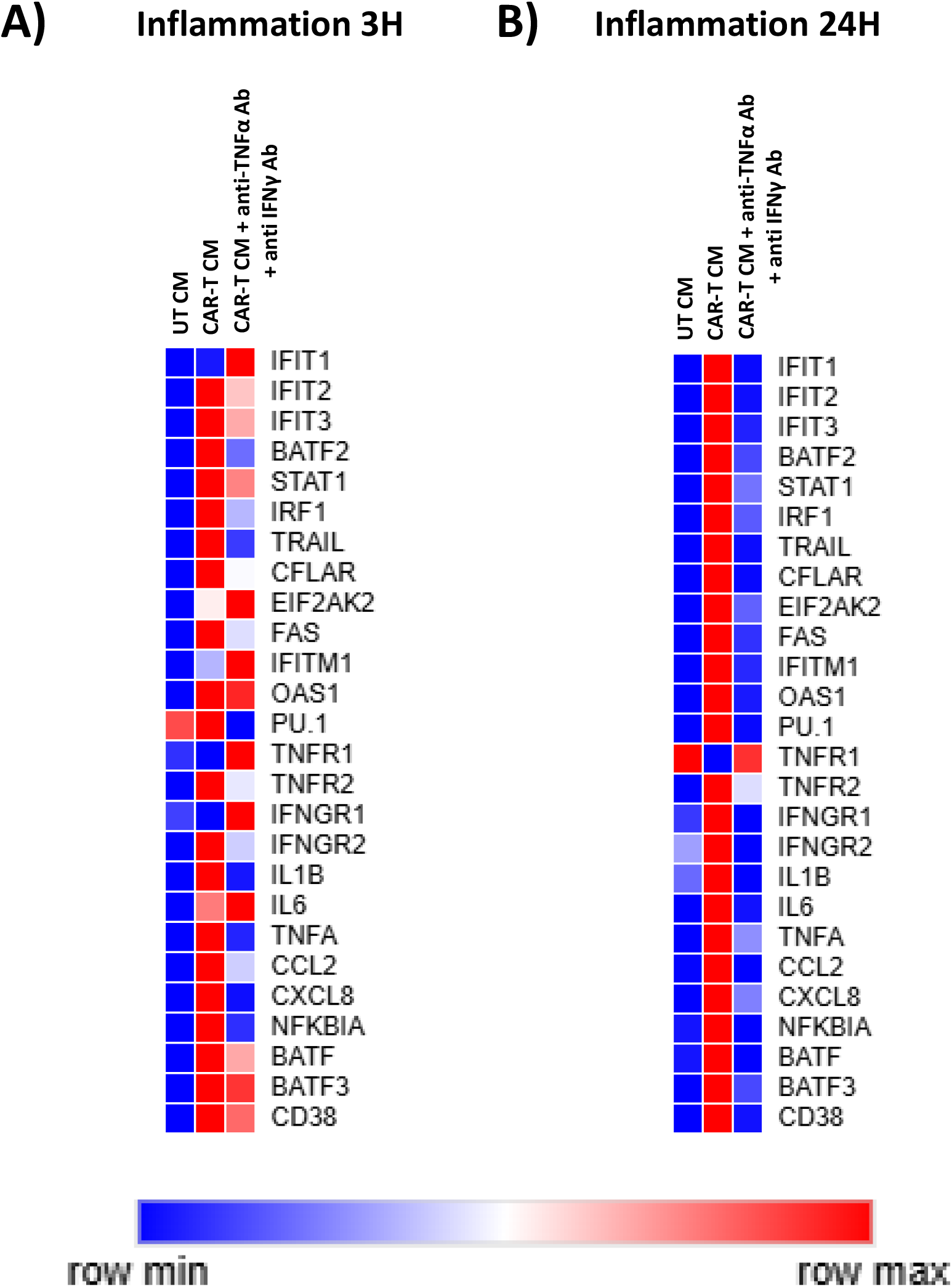

## References

1. June, C. H. & Sadelain, M. Chimeric Antigen Receptor Therapy. New England Journal of Medicine 379, 64–73 (2018).

2. Fried, S. et al. Early and late hematologic toxicity following CD19 CAR-T cells. Bone Marrow Transplant. 54, 1643–1650 (2019).

3. Rejeski, K., Hill, J. A., Dahiya, S. & Jain, M. D. Noncanonical and mortality-defining toxicities of CAR T cell therapy. Nature Medicine vol. 31 2132–2146 Preprint at 10.1038/s41591-025-03813-5 (2025).

4. Beyar-Katz, O., Rejeski, K. & Shouval, R. Immune effector cell-associated hematotoxicity: mechanisms, clinical manifestations, and management strategies. Haematologica vol. 110 1254–1268 Preprint at 10.3324/haematol.2024.286027 (2025).

5. Neelapu, S. S. et al. Chimeric antigen receptor T-cell therapy — assessment and management of toxicities. Nature Reviews Clinical Oncology 2017 15:1 15, 47–62 (2017).

6. Rejeski, K. et al. Immune effector cell–associated hematotoxicity: EHA/EBMT consensus grading and best practice recommendations. Blood 142, 865–877 (2023).

7. Caiado, F., Pietras, E. M. & Manz, M. G. Inflammation as a regulator of hematopoietic stem cell function in disease, aging, and clonal selection. J. Exp. Med. 218, (2021).

8. Dybedal, I., Bryder, D., Fossum, A., Rusten, L. S. & Jacobsen, S. E. W. Tumor necrosis factor (TNF)–mediated activation of the p55 TNF receptor negatively regulates maintenance of cycling reconstituting human hematopoietic stem cells. Blood 98, 1782–1791 (2001).

9. Yang, L. et al. IFN-γ Negatively Modulates Self-Renewal of Repopulating Human Hemopoietic Stem Cells. The Journal of Immunology 174, 752–757 (2005).

10. Strati, P. et al. Prolonged cytopenia following CD19 CAR T cell therapy is linked with bone marrow infiltration of clonally expanded IFNγ-expressing CD8 T cells. Cell Rep. Med. 4, (2023).

11. Khelil, M. Ben et al. CAR T cell–mediated bone marrow inflammation causes hematotoxicity and favors clonal hematopoiesis. Sci. Transl. Med. 17, 9790 (2025).

12. Murtadha, M. et al. A CD38-directed, single-chain T-cell engager targets leukemia stem cells through IFN-γ–induced CD38 expression. Blood 143, 1599–1615 (2024).

13. Juluri, K. R. et al. Severe cytokine release syndrome is associated with hematologic toxicity following CD19 CAR T-cell therapy. Blood Adv. 6, 2055–2068 (2022).

14. Read, J. A., Rouce, R. H., Mo, F., Mamonkin, M. & King, K. Y. Apoptosis of Hematopoietic Stem Cells Contributes to Bone Marrow Suppression Following Chimeric Antigen Receptor T Cell Therapy. Transplant. Cell. Ther. 29, 165.e1-165.e7 (2023).

15. Matatall, K. A. et al. Chronic Infection Depletes Hematopoietic Stem Cells through Stress-Induced Terminal Differentiation. Cell Rep. 17, 2584–2595 (2016).

16. Bailey, S. R. et al. Blockade or Deletion of IFNγ Reduces Macrophage Activation without Compromising CAR T-cell Function in Hematologic Malignancies. Blood Cancer Discov. 3, 136–153 (2022).

17. Manni, S. et al. Neutralizing IFNγ improves safety without compromising efficacy of CAR-T cell therapy in B-cell malignancies. Nature Communications 2023 14:1 14, 3423-(2023).

