## Supplemental methods and figure legends for "Sustained exposure to CAR-T cell secretome impairs human Hematopoietic Stem Cell function and is reversible by dual TNFα-IFNγ blockade"

### Materials and Methods

#### CAR-T cell production and condition medium generation

Human peripheral blood mononuclear cells (PBMCs) were isolated by density gradient centrifugation and activated with 50 ng/mL OKT3 in IL-2–containing T-cell medium. A T-cell medium for activation and transduction included RPMI 1640 supplemented with 10% FBS, 2 mmol/L L-Glu, 100 U/mL penicillin, 100 µg/mL Streptomycin, and IL2 (100 IU/mL). Activated T cells were transduced with vectors encoding CD19-targeting CAR constructs. Two CD19 CAR designs were used in this study: (i) an FMC63-based CAR containing a CD28-CD3ζ intracellular signaling domain cloned into a retroviral MSGV backbone[1], and (ii) a CD19 CAR containing a 4-1BB-CD3ζ signaling domain delivered using lentiviral construct[2]. Transduced cells were expanded for 7–11 days prior to use. CAR expression was confirmed by flow cytometry before functional experiments. NALM6 target cells (ATCC, CRL-3273) were grown in standard culture conditions, using RPMI medium supplemented with 10% fetal bovine serum (FBS), 2 mM L-Glutamate, 100 U/mL penicillin and 100 µg/mL streptomycin.

For generation of conditioned media (CM), CD19 CAR-T cells or donor-matched untransduced T cells were co-cultured with CD19+ NALM6 B-ALL target cells at 1:1 ratio for 48 hours. Following incubation, supernatants were collected, centrifuged to remove cellular debris, and stored at -80$^{o}$C until use. For HSPC exposure experiments, conditioned media were diluted 1:10 in cytokine-supplemented StemSpan SFEM medium (detailed below) prior to addition to CD34+ cells. Conditioned media generated from CD28-based and 4-1BB-based CD19 CAR-T cells produced comparable levels of IFNγ and TNFα as measured by ELISA and induced similar HSPC phenotypic and functional responses. Therefore, data derived from both constructs were analysed collectively.

#### CD34+ HSPCs purification

Cord blood units were obtained according to procedures approved by the institutional review boards of Tel Aviv University. Informed consent was obtained from all subjects. Two-six deidentified cord blood units were pooled, and mononuclear cells were isolated by density gradient centrifugation. CD34+ cells from cord blood or from the bone marrow samples were enriched by positive selection with MACS CD34+ ultra-pure kit (Miltenyi Biotech, Cat# 130-100-453) according to the manufacturer's instructions. Purified cells were stored in liquid nitrogen and thawed for subsequent experiments.

**CD34+ HSPC culture and conditioned medium exposure**

Thawed CD34+ cells were cultured in StemSpan SFEM II serum-free medium (Stem Cell Technologies, Cat# 09655) supplemented with recombinant human cytokines (PeproTech): SCF (25 ng/ml), FLT3L (25 ng/ml), and TPO (25 ng/ml).

CD34+ cells were cultured in UT CM or CAR-T CM (diluted 1:10) for 1, 3, or 7 days. For pulse experiments, cells were exposed to CAR-T CM for 3 hours, washed, and subsequently cultured in cytokine-supplemented StemSpan medium for the remainder of the culture period. Neutralizing antibodies against human IFNγ (clone- B27, Biolegend - 10ug/ml) and TNFα (Etanercept, MedChem Express – 10ug/ml) and the IL6 receptor antagonist LMT-28 (MedChem Express – 10uM) were added at the start of culture and maintained throughout 7-day exposure unless otherwise indicated.

**Flow cytometry**

Cells were stained with fluorochrome-conjugated antibodies against human CD34, CD38, CD45RA, EPCR, CD33, and CD11b. HSCs were defined as EPCR+CD34+CD33+CD45RA−, and HSPCs as CD34+CD33+CD45RA−. Data were acquired on Cytoflex4L or Cytoflex 5L flow cytometers and analysed using Cytexpert or FlowJo software (BD Biosciences).

**Immunodeficient mice repopulating cell assay**

All animal experimental protocols were approved by the Institutional Animal Care and Use Committee of Tel-Aviv University, Israel (TAU-MD-IL-2307-152-5). Mice were housed within the Tel Aviv University Specific Pathogen Free (SPF) facility in individually ventilated cages with four to five animals of the same sex per cage. All mice were maintained on a regular diurnal lighting cycle (12:12 light: dark) with ad libitum access to food and water.

For primary transplantation, CD34+ cells exposed ex vivo to UT CM or CAR-T CM for 1, 3, or 7 days were transplanted intravenously into female NSGW41 mice (8–12 weeks old) or busulfan-conditioned NSG mice (25 mg/kg busulfan intraperitoneally 24 hours prior to transplantation), as indicated. Primary and secondary transplants were performed in NSGW41 mice for the 1-day and 7-day exposure groups and in busulfan-conditioned NSG mice for the 3-day exposure group. For the 1-day and 3-day exposure groups, progeny of 50,000 CD34+ cells were transplanted per mouse. For the 7-day exposure group, progeny derived from 10,000 CD34+ cells plated at day 0 were transplanted. Mice were anesthetized with isoflurane (3% in 2.5% oxygen via a calibrated vaporizer), and a drop of ophthalmic lidocaine was applied to the eye immediately before injection. Cells were delivered intravenously into the retrobulbar plexus while animals remained under continuous isoflurane anesthesia via a nose cone. To ensure humane endpoints, mice were monitored daily for health parameters according to the protocol approved by the Institutional Animal Care and Use Committee of Tel-Aviv University (TAU-MD-IL−2307−152–5). Changes in any of the following parameters were defined as a human endpoint: body weight loss of 20% or more between the measurements (monitored by laboratory balances), lethargy, ruffled fur, hind limb paralysis, hunched posture and labored breathing (all monitored by visual inspection). Upon meeting the humane endpoint criteria or twelve weeks after the injection, mice were euthanized by CO2 inhalation using a fill rate of 50% of the chamber volume/minute. CO2 flow was maintained for at least 2 min after breathing stops to ensure death. Euthanasia was confirmed by the absence of breathing and reflexes. No secondary method of euthanasia was performed. Bone Marrow, Spleen and Peripheral Blood engraftment was analyzed by flow cytometry using hCD45, hCD33, hCD19 and hCD3 antibodies. 100,000 bone marrow mononuclear cells harvested from primary recipients were cultured in StemSpan SFEM II medium supplemented with cytokines and human plasma (10%) for 24 hours to eliminate residual murine cells. Subsequently, viable cells were plated in methylcellulose-based medium (Stem Cell Technologies, Cat# H4435), and colonies were enumerated after 14 days of incubation.

For secondary transplantation, 90% of whole bone marrow of primary mice (> 1% hCD45 chimerism) was reinjected into secondary NSGW41 mice or busulfan conditioned NSG mice intravenously into the retrobulbar plexus and the BM engraftment was analyzed by immunostaining and flow cytometry after additional 12 weeks.

**ELISA**

Cell free supernatants were collected from co-cultures and ELISA for the detection of human IFN-γ and TNF-α was carried out using Elisa MAX$^{TM}$ Deluxe kit (Biolegend) according to manufacturer's instructions.

**qRT-PCR analysis**

Total RNA was extracted using the RNeasy Micro kit (Qiagen Cat# 74004). cDNA synthesis was performed using the qScript cDNA Synthesis kit (Quanta Bio Cat# 95047-025). Real-time quantitative PCR was performed using PerfeCTa SYBR green supermix reagent (Quanta Bio Cat# 95054-500) and analysed by QuantStudio 5 Real-time PCR system (ThermoFisher). Relative expression was calculated for each gene using by $2^{-\Delta\Delta CT}$ method. *GAPDH* was used for normalization.

**CAR-T killing assay**

CAR-T cells were co-cultured with NALM6 target cells in the presence or absence of cytokine blockade. After 24 hours, cells were analysed by flow cytometry, and proportions of CD3+ and CD19+ cells were quantified.

**Statistical analysis**

Statistical analysis was performed with GraphPad Prism 9, using student's t-test. Statistical significance is defined as $p<0.05$. Bars represent Mean ± Standard error of mean (SEM) of independent experiments. *P≤0.05, **P≤0.01, ***P≤0.001, ****P≤0.0001.

**Supplementary Figure 1.** ***In vivo*** **functional consequences of CAR-T conditioned medium exposure**

**(A)** Primary peripheral blood hCD45+ levels at 12 weeks following transplantation of CD34+ cells exposed for 1, 3, or 7 days. **(B)** Primary spleen hCD45+ levels at 12 weeks. **(C)** Secondary peripheral blood engraftment at 12 weeks. **(D)** Secondary spleen engraftment at 12 weeks. Individual mice and median values are shown.

**Supplementary Figure 2. Acute CAR-T conditioned medium induces IFNγ-dependent CD38 upregulation in human HSPCs**

**(A)** Representative flow cytometry plots of CD34+ HSPCs cultured for 24 hours in UT CM or CAR-T CM, showing CD38 expression and EPCR+ fraction within the CD34+ compartment. **(B)** Quantification of CD38 median fluorescence intensity (MFI) in CD34+CD45RA- HSPCs cultured for 24 hours in UT CM or CAR-T CM in the presence of isotype control or IFNγ-neutralizing antibody. **(C)** Fold change in CD38 MFI relative to UT CM controls under the indicated conditions. **(D)** CD38 MFI in CD34+CD45RA- HSPCs cultured with recombinant cytokines for 24 hours. **(E)** Fold change in CD38 MFI relative to untreated controls following recombinant cytokine exposure. Data are presented as mean ± SEM from independent experiments. Statistical analysis was performed using unpaired Welch's t test or one-way ANOVA as appropriate.

**Supplementary Figure 3. Sustained CAR-T conditioned medium drives impaired HSPC expansion and myeloid skewing**

**(A)** Frequency of EPCR+CD34+CD45RA− HSCs after 24-hour exposure to UT CM or CAR-T CM. **(B)** Fold change in frequency of EPCR+CD34+CD45RA− HSCs after 24-hour exposure to UT-CM or CAR-T-CM, normalized to UT-CM controls. **(C)** Representative flow cytometry plots of CD34+ HSPCs after 7-day culture under indicated conditions. **(D–G)** Absolute expansion of sorted EPCR+CD34+CD38−CD45RA− HSCs cultured for 7 days in UT CM or

CAR-T CM, showing (A) HSC numbers, (B) HSPC numbers, (C) CD11b+ myeloid cell numbers, and (D) CD38 MFI in CD34+CD45RA- HSPCs. **(H)** Fold change in CD38 MFI relative to UT CM controls. Data are presented as mean ± SEM. Statistical analysis was performed using unpaired Welch's t test or one-way ANOVA as appropriate.

**Supplementary Figure 4. CAR-T conditioned medium impairs HSPC expansion independent of apoptosis or necroptosis**

**(A–B)** Absolute expansion numbers (A) and fold change (B) of EPCR+CD34+CD45RA− HSCs after 7-day culture. **(C–D)** Absolute expansion (C) and fold change (D) of CD34+CD33+CD45RA− HSPCs after 7-day culture. **(E–F)** Absolute expansion (E) and fold change (F) of CD34−CD33+CD11b+ myeloid cells after 7-day culture. **(G–H)** CD38 median fluorescence intensity (MFI) in HSPCs (G) and corresponding fold change relative to UT CM controls (H) after 7-day culture. **(I-J)** Absolute expansion of EPCR+CD34+CD33+CD45RA− HSCs (I) and fold change (J) and CD34+CD33+CD45RA- HSPC numbers (K) and fold change (L) after 7-day culture in CAR-T CM in the presence of Q-VD (caspase inhibitor), Nec-1s (necroptosis inhibitor) and MCC950 (NLRP3 inhibitor). Data are presented as mean ± SEM.

**Supplementary Figure 5. Cytokine blockade partially rescues primitive HSC phenotypes**

**(A–B)** Absolute expansion (A) and fold change (B) of EPCR+CD34+CD45RA− HSCs after 7-day culture in CAR-T CM with indicated cytokine inhibitors. **(C–D)** Absolute expansion of CD34+CD33+CD45RA− HSPCs (C) and CD34−CD33+CD11b+ myeloid cells (D). **(E–F)** CD38 MFI (E) and fold change (F) in CD34+ HSPCs after 7-day exposure to CAR-T CM with cytokine blockade. Data are presented as mean ± SEM.

**Supplementary Figure 6. Functional rescue and CAR-T cytotoxicity**

**(A–B)** Primary peripheral blood (A) and spleen (B) hCD45+ engraftment at 12 weeks following 7-day CAR-T CM exposure ± combined IFNγ and TNFα blockade. **(C–D)** Secondary peripheral blood (C) and spleen (D) engraftment. **(E)** CAR-T cytotoxicity assay after 24-hour co-culture with NALM6 target cells in the presence of indicated cytokine inhibitors. Proportions of CD3+ and CD19+ cells are shown. Individual mice and median values are shown for in vivo experiments.

**Supplementary Figure 7. Cytokine blockade reverses inflammatory transcriptional programs**

**(A–B)** Heatmaps of qRT-PCR analysis of CD34+ HSPCs treated with UT CM, CAR-T CM, or CAR-T CM with combined IFNγ and TNFα blockade for 3 hours (A) or 24 hours (B). Data represent mean expression from three independent biological experiments with technical replicates.

### Supplemental references


1. Rozenbaum, M. *et al.* Genotoxicity Associated with Retroviral CA R Transduction of ATM-Deficient T Cells. *Blood Cancer Discov.* **5**, 267–275 (2024).
2. Carnevale, J. *et al.* RASA2 ablation in T cells boosts antigen sensitivity and long-term function. *Nature* **609**, 174–182 (2022).

| Antigen | Clone | Fluorophore | Dilution | Manufacturer | Catalog number |
|---|---|---|---|---|---|
| CD34 | 581 | FITC | 1-100 | Biolegend | 343504 |
| | 581 | PE | 1-100 | Beckman Coulter | A07776 |
| | 8G12 | APC | 1-100 | BD Bioscience | 345804 |
| CD38 | HB7 | PC7 | 1-100 | Biolegend | 356608 |
| CD45RA | HI100 | BV605 | 1-200 | Biolegend | 304134 |
| CD201 | RCR-401 | PE | 1-100 | Biolegend | 351904 |
| CD33 | P67.6 | PE | 1-100 | Biolegend | 366608 |
| CD19 | SJ25C1 | APC | 1-100 | Biolegend | 363006 |
| CD11b | M1/70 | PC5.5 | 1-100 | Biolegend | 101228 |
| CD3 | REA613 | FITC | 1-100 | Miltenyi Biotec | 130-114-138 |
| CD45 | J33 | PC7 | 1-100 | Beckman Coulter | IM3548 |
| Ultra-LEAF™ Purified Mouse IgG1, κ Isotype Ctrl Antibody | MGI-45 | | 10ug/ml | Biolegend | 401407 |
| Ultra-LEAF™ Purified anti-human IFN-γ | B27 | | 10ug/ml | Biolegend | 506532 |
| Etanercept | | | 10ug/ml | MedChem express | HY-108847 |
| LMT-28 | | | 10uM | MedChem express | HY-102084 |
| Stem Cell Factor (SCF) | | | | Peprotech | 300-07-100UG |

| | | | | | |
|---|---|---|---|---|---|
| FLT3 ligand (FLT3) | | | | Peprotech | 300-19-100UG |
| Thrombopoietin (TPO) | | | | Peprotech | 300-18-50UG |
| ELISA MAX™ Deluxe Set Human IFN-γ | | | | Biolegend | 430115 |
| ELISA MAX™ Deluxe Set Human TNF-α | | | | Biolegend | 430215 |

**List of primers**

| Gene name | Forward | Reverse |
|---|---|---|
| GAPDH | TTC GTC ATG GGT GTG AAC CA | CTG TGG TCA TGA GTC CTT CCA |
| IFIT1 | GGA CCC TGA AAA CCC TGA AT | TGT GGC TAA TTT AAA GCC ATC C |
| IFIT2 | GCA AGC TAC CGT CTG GAC A | CTT GCC TCA GAG GGT CAA TG |
| IFIT3 | GTG CTG CTA CAA GGC AAA AGT | TCA TCT CTT TAT TTC CAC TAG CTT CA |
| BATF | GAT GTG AGA AGA GTT CAG AGG AG | GTT TCT CCA GGT CTT CGC TCT C |
| BATF2 | GCT GAA GAA GCA GAA GAA CCG G | TGC AGG GAC TGG ATC TCC TTC C |
| BATF3 | ACC GAG TTG CTG CTC AGA GAA G | AGG TGC TTC AGC TCC TCT GTC A |
| STAT1 | CCT GCT GCG GTT CAG TGA | GGT TCA ACC GCA TGG AAC TC |
| IRF1 | GAG GAG GTG AAA GAC CAG AGC A | TAG CAT CTC GGC TGG ACT TCG A |
| TRAIL | CCA AAA GTG GCA TTG CTT GTT | ACG GAG TTG CCA CTT GAC TTG |
| CFLAR | CTC TTT TTG TGC CGG GAT GT | AAG TCC CCG ACA GAC AGC TTA C |
| EIF2AK2 | GGC ATT CAG CTC CAC ACT TG | ACA GAC GAG TGA TAC CAG CG |
| FAS | TTG GTG GAC CCG CTC AGT A | AGC AAT CCT CCG AAG TGA AAG A |
| IFITM1 | AAG CCA GAA GAT GCA CAA GGA | GGA GGT CTC GCT GTG GAT GT |
| OAS1 | CGAGGGAGCATGAAAACACATTT | GCAGAGTTGCTGGTAGTTTATGAC |
| SPI1 | GAC ACG GAT CTA TAC CAA CGC C | CCG TGA AGT TGT TCT CGG CGA A |
| IFNGR1 | AGT GCT TAG CCT GGT ATT CAT CTG | GGC TGG TAT GAC GTG ATG AGT G |

| | | |
|---|---|---|
| IFNGR2 | CTC CAT TCT GCC TGG GTG ACA A | CGT GGA GGT ATC AGC GAT GTC A |
| TNFRSF1A | TGT GTC TCC TGT AGT AAC TGT AAG | AGT CCT CAG TGC CCT TAA CA |
| TNFRSF1B | GTC CAC ACG ATC CCA ACA C | CAC ACC CAC AAT CAG TCC AA |
| IL1B | CCACAGACCTTCCAGGAGAATG | GTGCAGTTCAGTGATCGTACAGG |
| IL6 | AGACAGCCACTCACCTCTTCAG | TTCTGCCAGTGCCTCTTTGCTG |
| TNFA | CTCTTCTGCCTGCTGCACTTTG | ATGGGCTACAGGCTTGTCACTC |
| CCL2 | AGAATCACCAGCAGCAAGTGTCC | TCCTGAACCCACTTCTGCTTGG |
| CXCL8 | GAGAGTGATTGAGAGTGGACCAC | CACAACCCTCTGCACCCAGTTT |
| NFKBIA | TCCACTCCATCCTGAAGGCTAC | CAAGGACACCAAAAGCTCCACG |
| CD38 | TCT TGC CCA GAC TGG AGA AAG G | TGG ACC ACA TCA CAG GCA GCT T |
